# SoundChunk: A free open-source and user oriented R package for acoustic data management, sound detection and extraction

**DOI:** 10.1101/2023.08.29.555312

**Authors:** Avelyne S. Villain, Paul Renaud-Goud

## Abstract

In a standard bioacoustic experiment setting, after data collection and before data analysis lies a time-consuming process that consists in extracting and labeling sounds of interest (chunks) from usually long recording files, collected through spreading PAM (*Passive Acoustic Monitoring*) for instance, and storing them in a structured and exploitable way based on both the meta-data of the initial recordings and the label data.

`SoundChunk` is a user-oriented package in R that aims at providing tools for any R user so that they can go efficiently and comfortably through this process. Usually, the tasks include detecting, labeling, and extracting sound from audio files. With the recent advance of Machine Learning, especially Convolutional Neural Networks, some of these tasks are integrated in the ML framework; however, the learning phase relies on labeled chunks that need to be created.

Three types of data are involved in the process: (i) WAVE files, whether they come from the initial recordings or are generated by chunk extraction, (ii) meta-data describing the conditions in which the initial WAVE files were recorded, (iii) label track files, whether they are automatically generated or manually input, that contain timed information about audio files. `SoundChunk` provides utilities to sanely manipulate each of these three categories and combine them into easy-to-use chunks along with their meta-data.

We expose functionalities that: (i) clean and restructure label tracks that are readable by softwares like *Audacity*, (ii) chop long recordings into small fixed-size slices, (iii) detect chunks both interactively (so that robust detection settings can be found on a subset of the recordings) and automatically (so that the detection is applied across all recordings), (iii) dispatch chunks into structured folder(s), according to their meta-data.

**Maintainers:** The team is open for suggestions and contributions. Contact: (currently: ASV, PRG).

**Users:** This document is available as a vignette once the package is loaded an may be used on a set of example data (Villain and Renaud-Goud 2023).

## Introduction

### Purpose of the package

`SoundChunk` is user-oriented and help:

- detecting sounds of interest in audio file
- building modular databases once sounds of interests have been identified

This package does not provide a 100% accurate detection but allows users to check the detections, edit and clean a pre-processed labeled audio to then build modular databases. The user thus keeps control on its data extraction for further analyses.

This package is built using in R (R Core Team 2023) and uses `seewave` (Sueur, Aubin, and Simonis 2008) (v2.2.0) and `tuneR` (Ligges et al. 2023) (v1.4.4) packages. Detection of sound of interest is based on the `timer()` and `ffilter()` functions, thus working on the smooth envelop of a sound file within a frequency band of interest.

The package allows bridging *R* and *Audacity* (“Audacity®” 2017), a free open-source and easy-to-use software for viewing, labeling and editing audio files.

This workflow gives an overview of the environment of the package and how it can be used on a set of example data (Villain and Renaud-Goud 2023).

## Definitions

The package uses a lexical field and some definitions may be required to fully experience it.

### Terminology

`Chunk` – sound unit of interest, could be a vocalisation, a hearth beat, a series of woodpecker sounds, a tap dance sound… The chunk represents the final sound unit we aim at detecting, store and analyse.

`Slice` – an extract of a longer audio file. Detection of chunks may be time-consuming and it could be of interest to apply it on slices. They are usually of fixed size and overlap so that no chunk is missed.

`label_track` – name given to a text file (tab separated value), exported or not from *Audacity*, that contains at least three columns: `start`, `end`, `label` and as many rows as chunks or slices.

`Origin` – name of a file without its extension. Label tracks usually share the same origin with their associated audio file.

### Actions

`Chunking` – the process of detecting and labeling chunks.

`Dispatching` – the process to extract chunks from an audio file based on their boundaries, which are stored in a label track (audacity-like `csv` files) and store them individually.

`Chopping` – the process of cutting a long recording into standardized pieces (whatever the recording contains).

## Main content of V1.0.0

1. Create databases from entire directories containing audio and label track files:
  - `dispatch_chunks()` : extract, name and save chunks locally
    - a `grouped` behaviour (one directory created per wanted groups of chunks),
    - a `flat` behaviour (all chunks from all recordings in a single directory)
  - to generate them, the package needs:
    - a `parameters` file with the match between the `origin` (*a*.*k*.*a*. the file names of the audio files without extension) and all common information associated to each recording. This file must be a tab separated (`.tsv`) file, with headers corresponding to the parameters associated to each recording. The `origin` column must be present.
    - a `naming_rule` file: containing information about the chunk and about the recording the user may want to see in the file name of each chunk and in the final log of the database. `set_naming_rule()`will help creating it and `set_naming_rule_path()` will allow using an already created `naming_rule` file.
2. Chunk a mono recording:
  - `detect_chunks_interactive()`: plot spectrograms and detections plots on audio files. Assist the user in finding *robust* parameters for chunking, according to the type of chunks. This program interactively asks the user:
    - if they want to change the detection settings and keep track of them,
    - to semi-manually label chunks to create semi-manual databases later on,
    - if they want some fragments of background noise to be labeled for further use.
  - `chop_recordings()`: when used along with `dispatch_chunks()`, generate slices of recordings for using `detect_chunks_interactive()`
  - `detect_chunks_auto()`: automatically detect all chunks in recordings, according to the chosen detection settings and normalisation value.
3. Interface between *Audacity* and *R*
  - `clean_audacity()`: clean label tracks exported from *Audacity* and create dataframes with headers and at least four columns: `start`, `end`, `label`, `index`
  - `expand_labels()`: modify the label track and add as many columns as type of information included in the label, based on a unique separator between pieces of information.
4. Note that all functions running on the entire content of a directory are also available as single file use.

## Installation and usage

The first time used, the package needs to be downloaded it from a *GitLab* server. Once it has been downloaded and locally stored, only loading the library will be needed to use the package. Install `SoundChunk`, using the `remotes` library

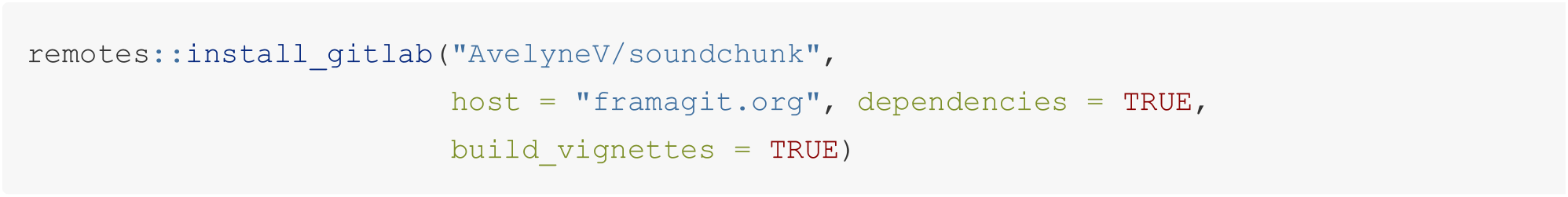

Use `SoundChunk`

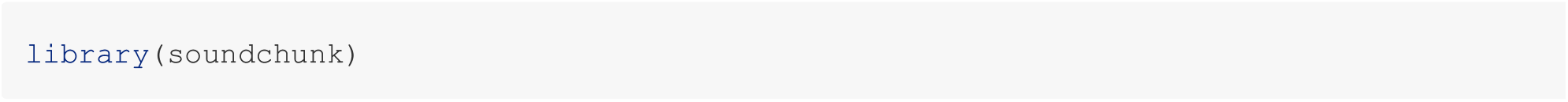

Information about `SoundChunk`, especially vignettes, or any of its function `?name_of_the_function`.

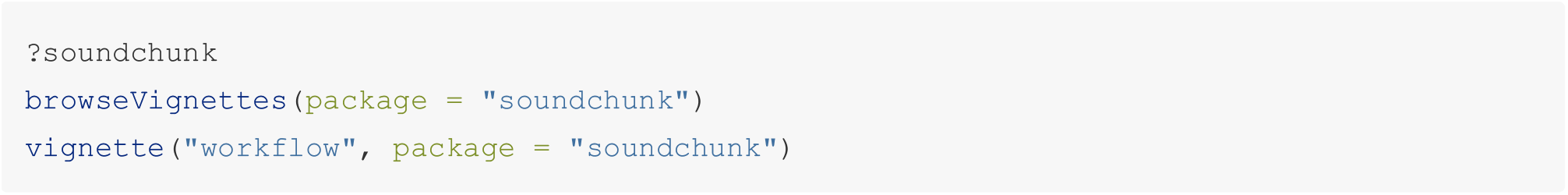

## Environment

The package comes with its environment as a global variable. We use this environment as a storage for settings that are widely used by the functions of the package. The status of these settings may be changed using a set of functions we will go through in the following sections. Note that if some status needs to be changed for your usage of the package, the user will have to change them every time the package is loaded.

### WAV_EXT

Only audio files in the `WAV` format are allowed. From one recorder to the other, different extensions may be used: either `.wav` or `.WAV`. The user may visualize this setting by calling `get_wav_ext()`

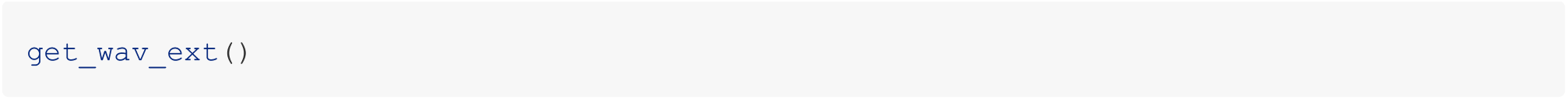

The default is`.WAV` however the user may change this feature to `.wav` using the `set_wav_ext()` function prior to using the functions of the package.

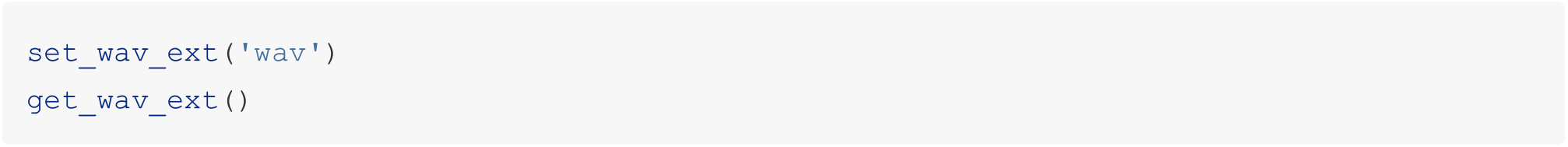

### FFT_WINDOW_LENGTH

When the function requires the computation of a Fast Fourier Transform, a window length has to be chosen. The size of this window mainly depends on the type of chunks.

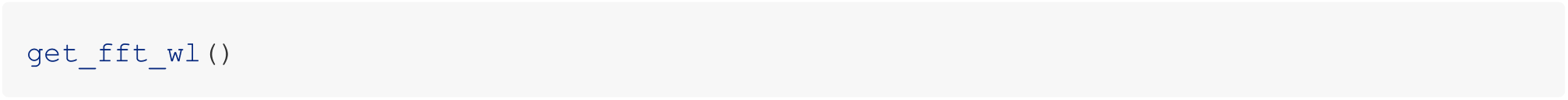

By default `FFT_WINDOW_LENGTH = 512` but it may be changed in the environment of the package using `set_fft_wl()`

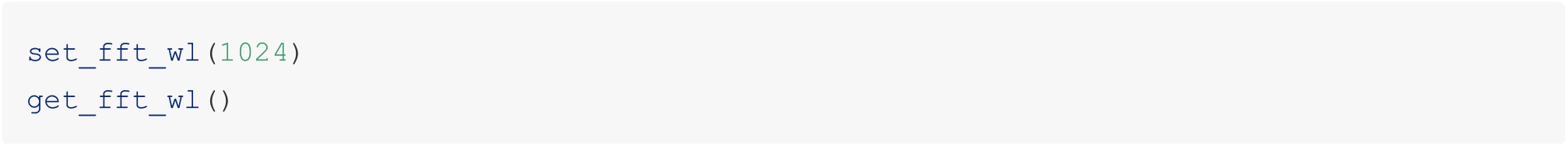

### AUDIO_PLAYER_PATH

The function `detect_chunks_interactive()` gives the **option** to listen to the audio piece. The variable `AUDIO_PLAYER_PATH` provides the path to the external software that is to be used. Default is `NULL`.

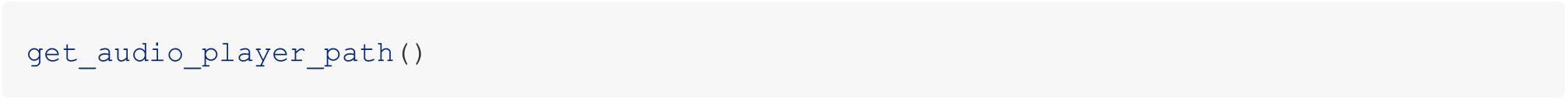

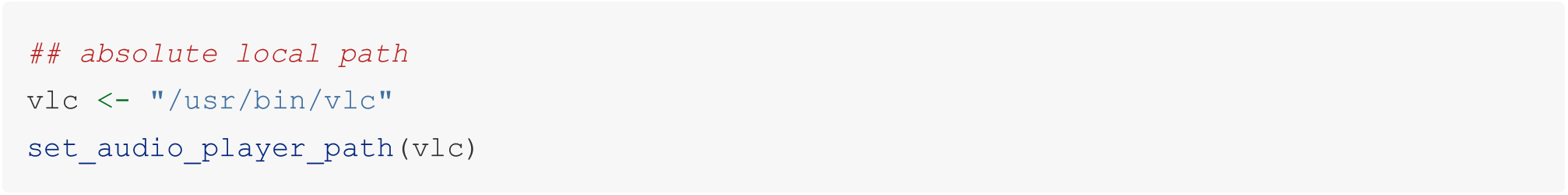

### NAMING_RULE and NAMING_RULE_PATH

In order to generate databases, a match between the context of a recording (*e*.*g*. date, field, treatment), the properties of a chunk (*e*.*g*. low bleat, tap) and the name of the chunk file is needed. Such a correspondence may be created interactively using `set_naming_rule()`. Once created, this information is stored into a global variable `NAMING_RULE` and stored in the file system, so that it may be called later using `set_naming_rule_path()`. By default, `NAMING_RULE` and `NAMING_RULE_PATH` are `NULL`.

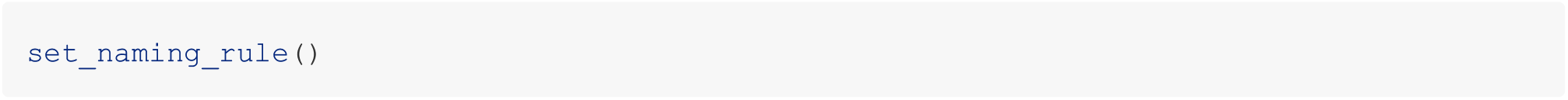

*N*.*B*. when creating a `naming_rule`, the name of the factor we input must match with the column names of the parameters file (type of information about the recording) and to the column names the label tracks (type of information about the chunk).

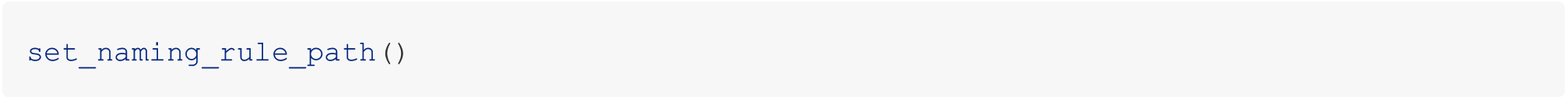

### SHOW_STEPS behaviour

For most of the functions contained in the package, the program may be paused to monitor the different steps of the program. The status of this variable may be changed using `show_steps()`. By default is `SHOW_STEP` is `FALSE`

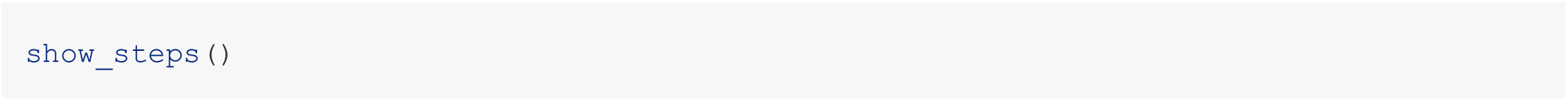

This status may be changed using `set_show_steps()`

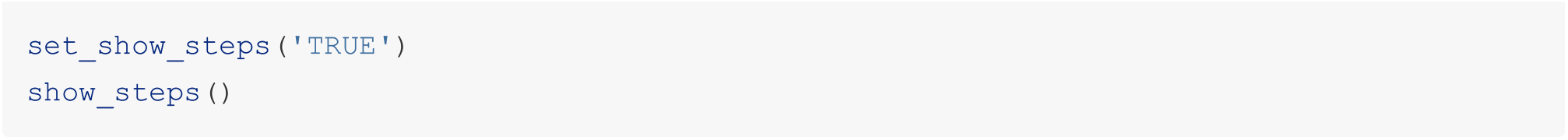

### Global variables for spectrogram settings

For spectrogram plots, some default values of the arguments of the `spectro()` function were chosen but may be edited.

`set_fast_display_spectro()`might especially be of interest to get faster plotted but lower quality spectrograms.

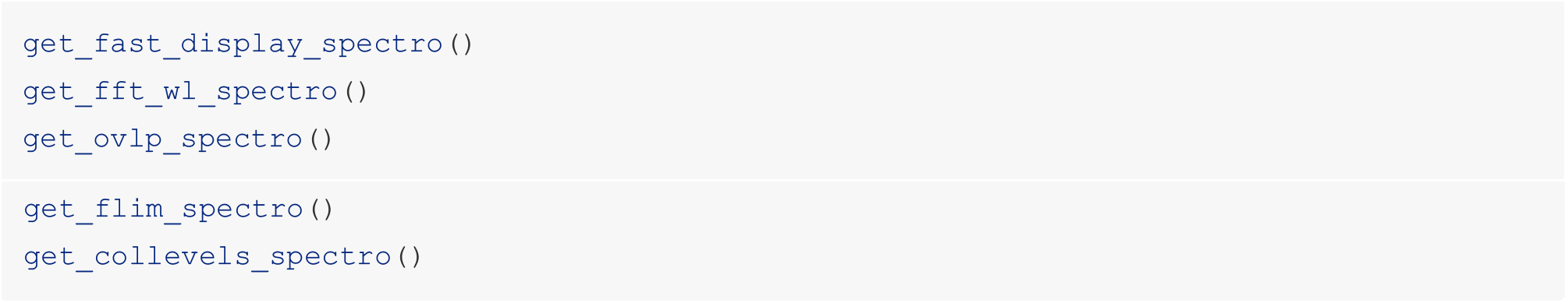

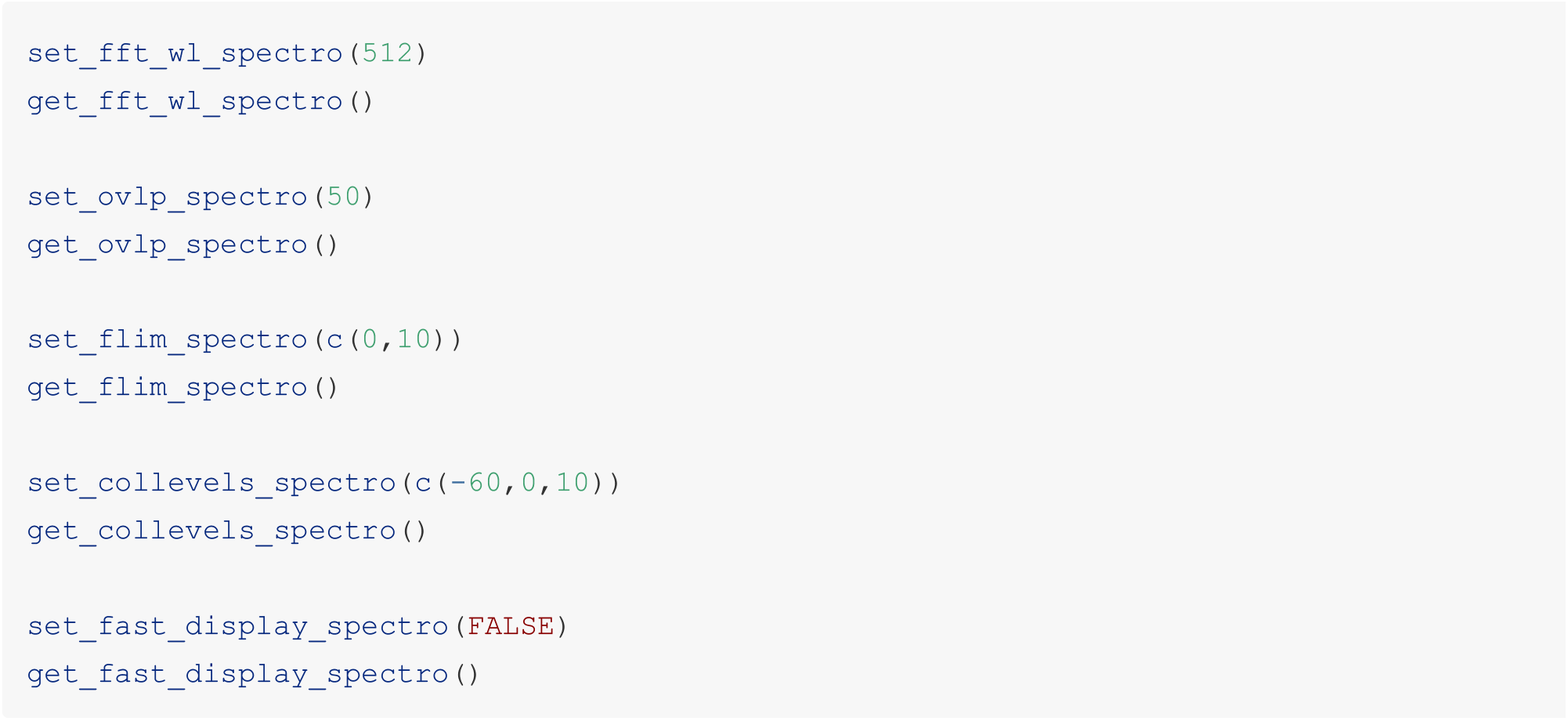

(Part of) the package environment may be visualised using `print_env_settings()`

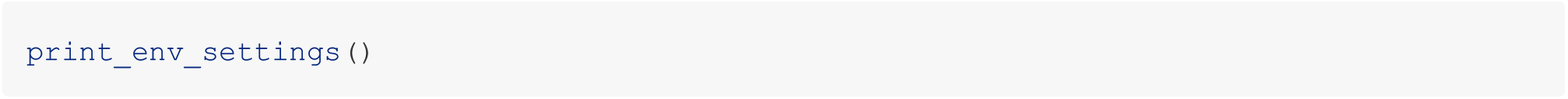

## Example workflows

In the following section, two scenarios are used to provide examples of how the package may be used:

- How to use `SoundChunk` to generate a modular database of labeled chunks,
- How to (semi)automatically detect chunks in audio files.

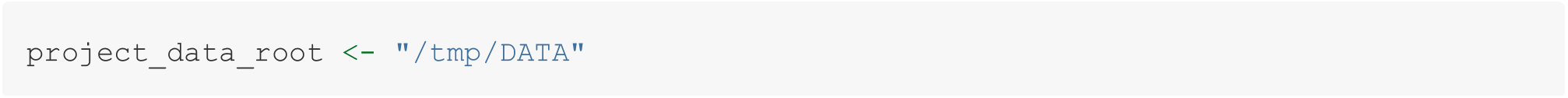

## 1 The labeled data scenario

In this scenario, we have labeled audio files, so we know when chunks occur and the information about each chunks is already stored in a label track.

Objective: the user may want to create databases, automatically and in the shape they desire! The package is beneficial in:

- finding the match between information of the recording contained in the parameters file and the audio file
- finding the information contained in the labels of the label track
- setting up a naming rule to choose what should be the name of the chunks and in the log of the chunk meta data
- dispatching the chunks into different types of databases
- creating log files to monitor the chunks and their information

### 1.1 Input data

- a bunch of audio files in a directory associated to each recording, a label track file whose name holds a different extension but the same name before extension. A label track contains three tab-separated columns: `start`, `end`, `label` of each chunk (could have been exported from *Audacity* for instance)
- a parameters file containing information about the recordings

What the directory looks like:

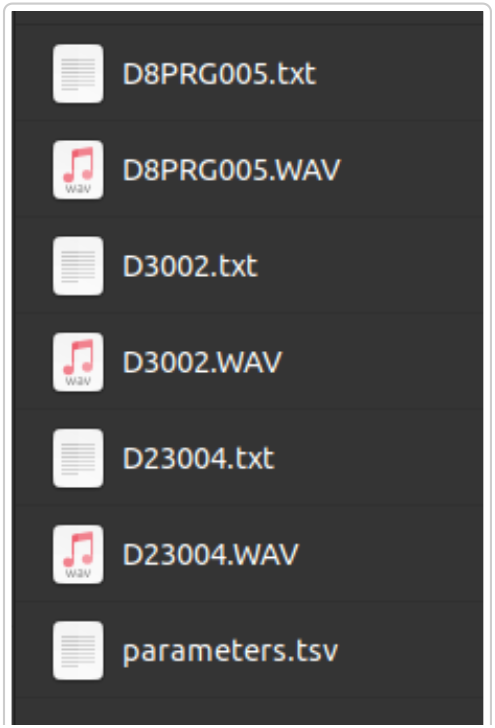

Architecture of input data

What the label track from *Audacity* looks like:

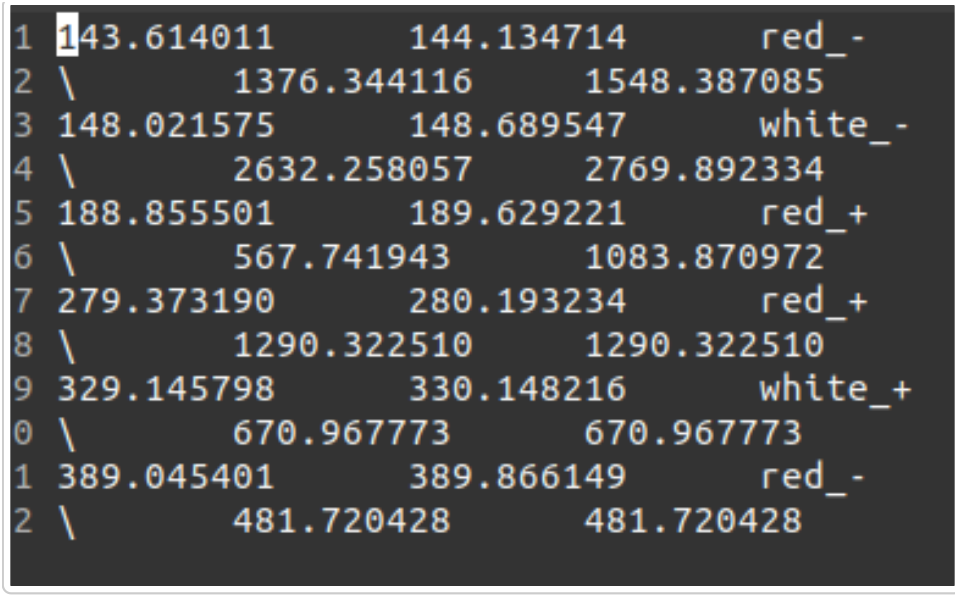

Input label track from Audacity

What a parameters file looks like.

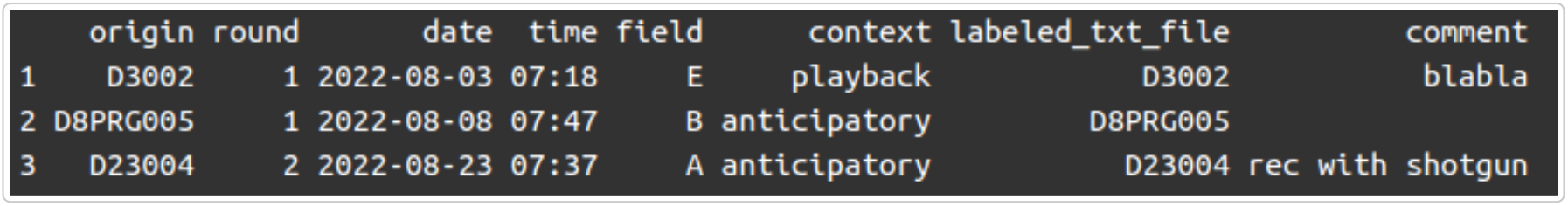

Parameters file. *Tab separated table with* .*tsv* *extension, with the column named* `origin` *with the name of the audio file, without extension*

### 1.2 Clean *Audacity* files

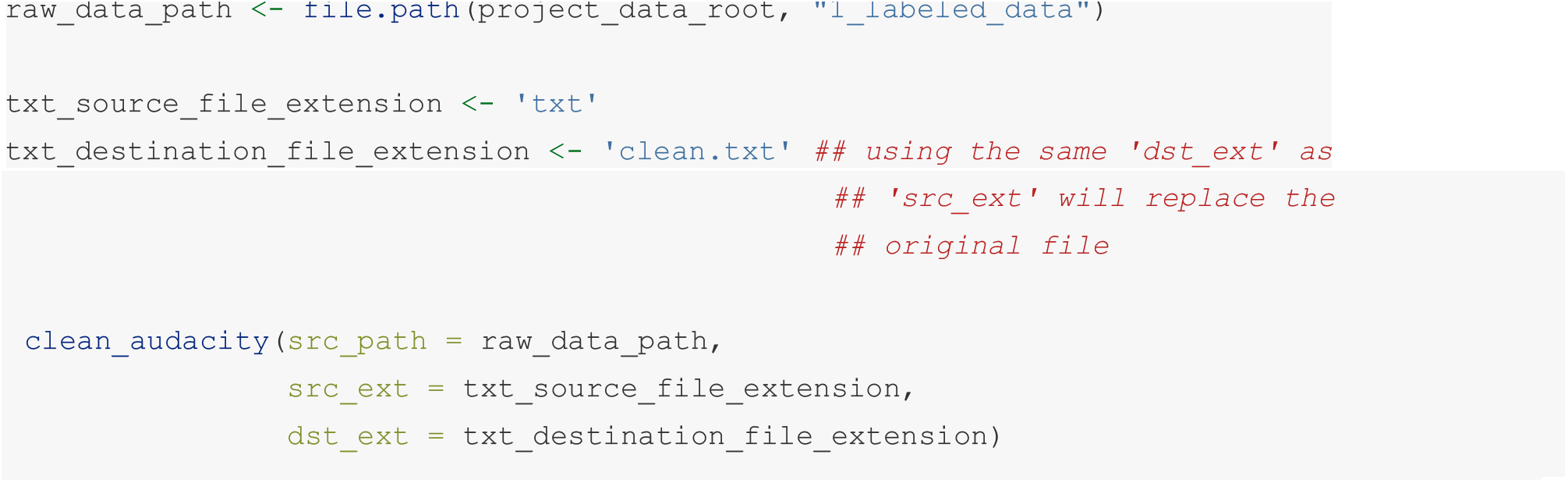

What the clean *Audacity* looks like:

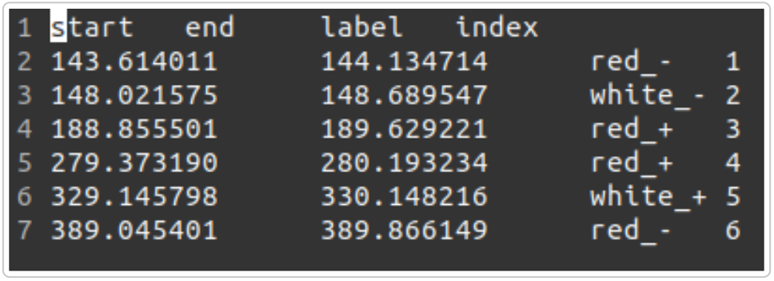

Cleaned label track

### 1.3 Expand label information

If the label contains several pieces of information, stored in a standard way, the user may extract them. Note that using space as a separator in the labels are not supported.

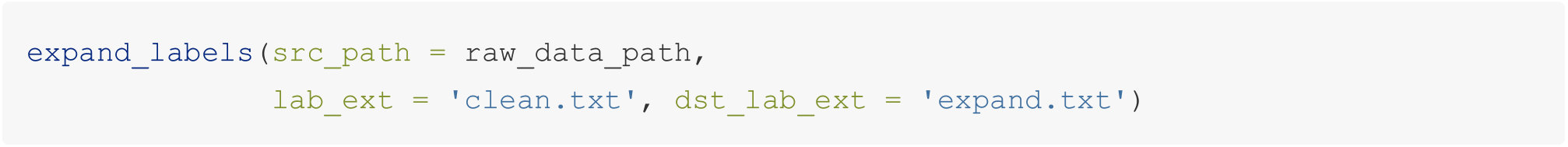

Label track contains `subject` and `quality` information.

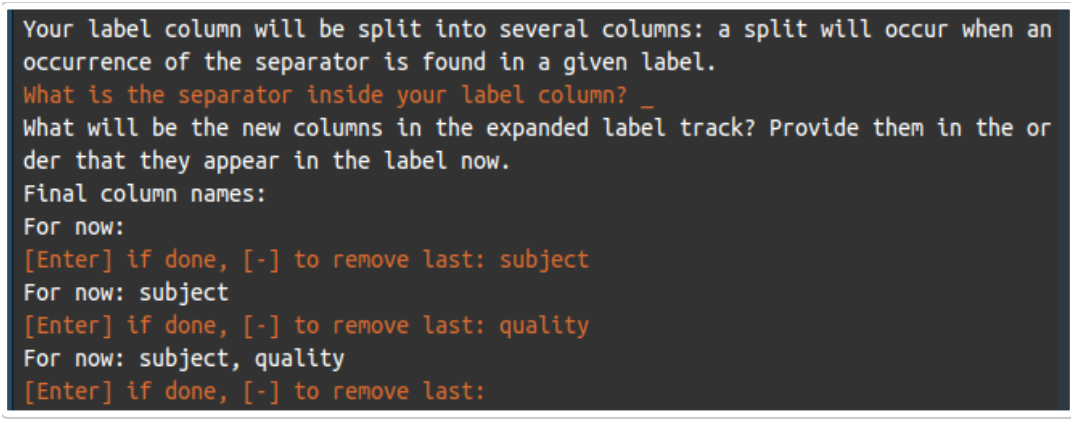

`expand_labels()`: user interaction

This is how the label track now looks like:

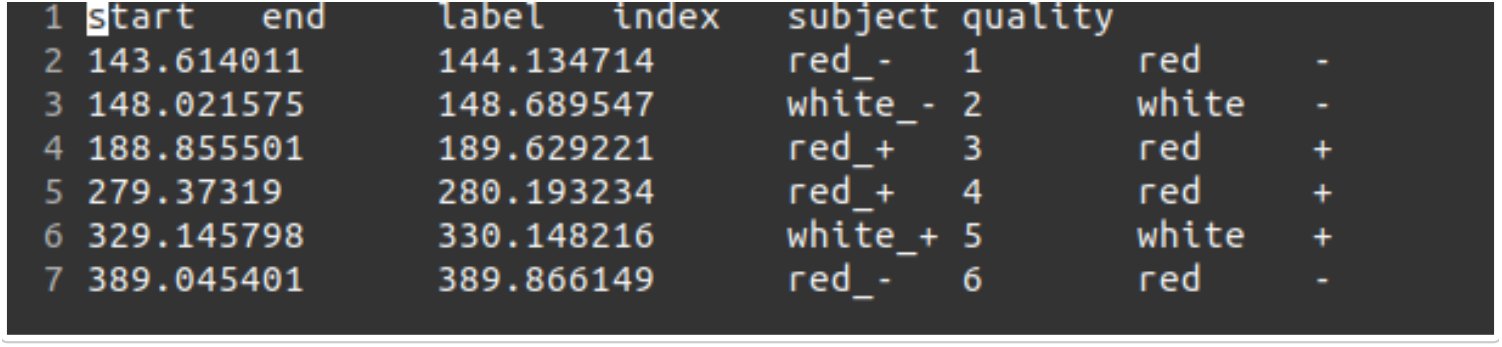

Expanded label track

### 1.4 Dispatch the chunks…

#### 1.4.1 …Into a flat database

Set up a `naming_rule`. A naming rule will allow us to choose the structure and content of the future file names of the database. It may be created interactively using `set_naming_rule()` or, if already created using `set_naming_rule_path()`.

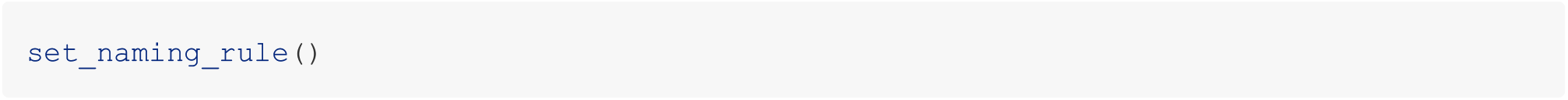

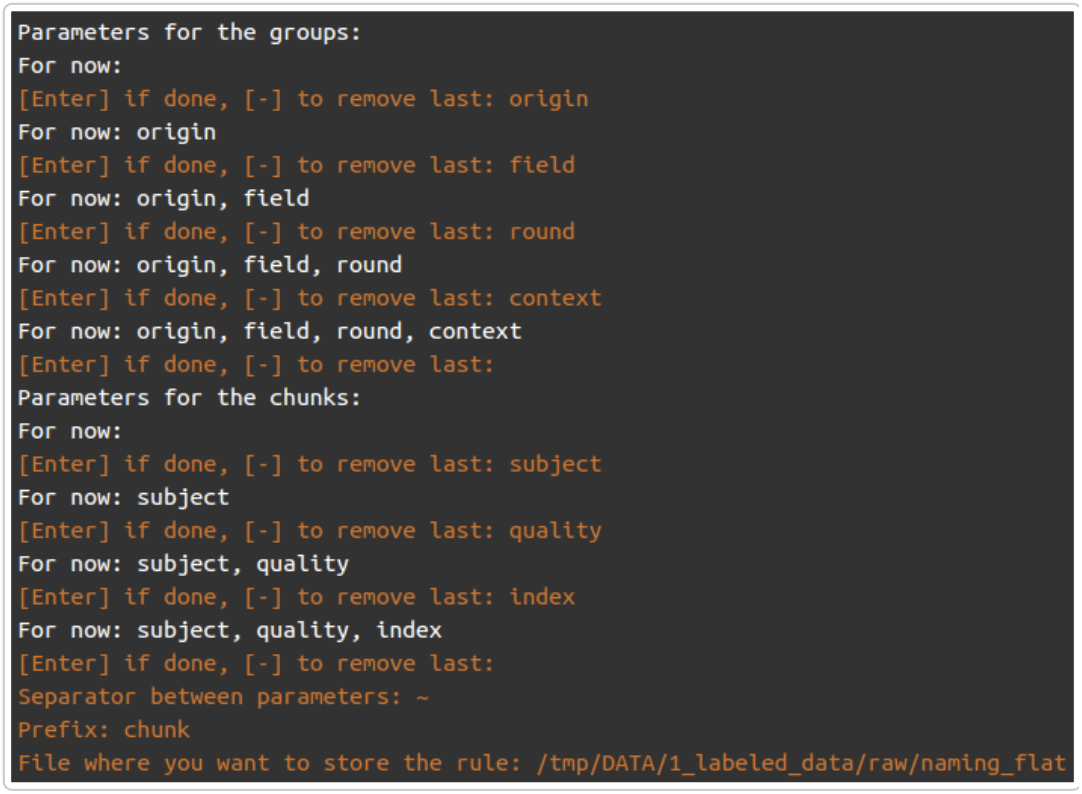

`set_naming_rule()`: user interaction

We have now assigned the global variable `NAMING_RULE`. If another one should be used, the global variable may be re assigned using `set_naming_rule_path()`.

What the naming rule called `naming_flat` looks like (through a call to print_env_settings()):

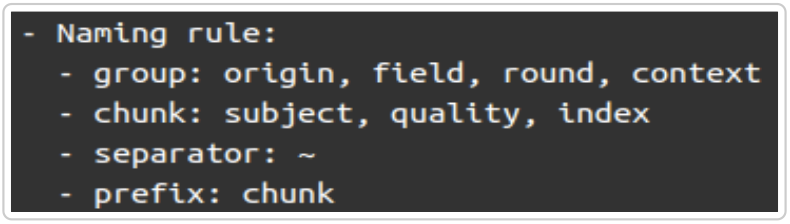

Naming rule for dispatching into flat database

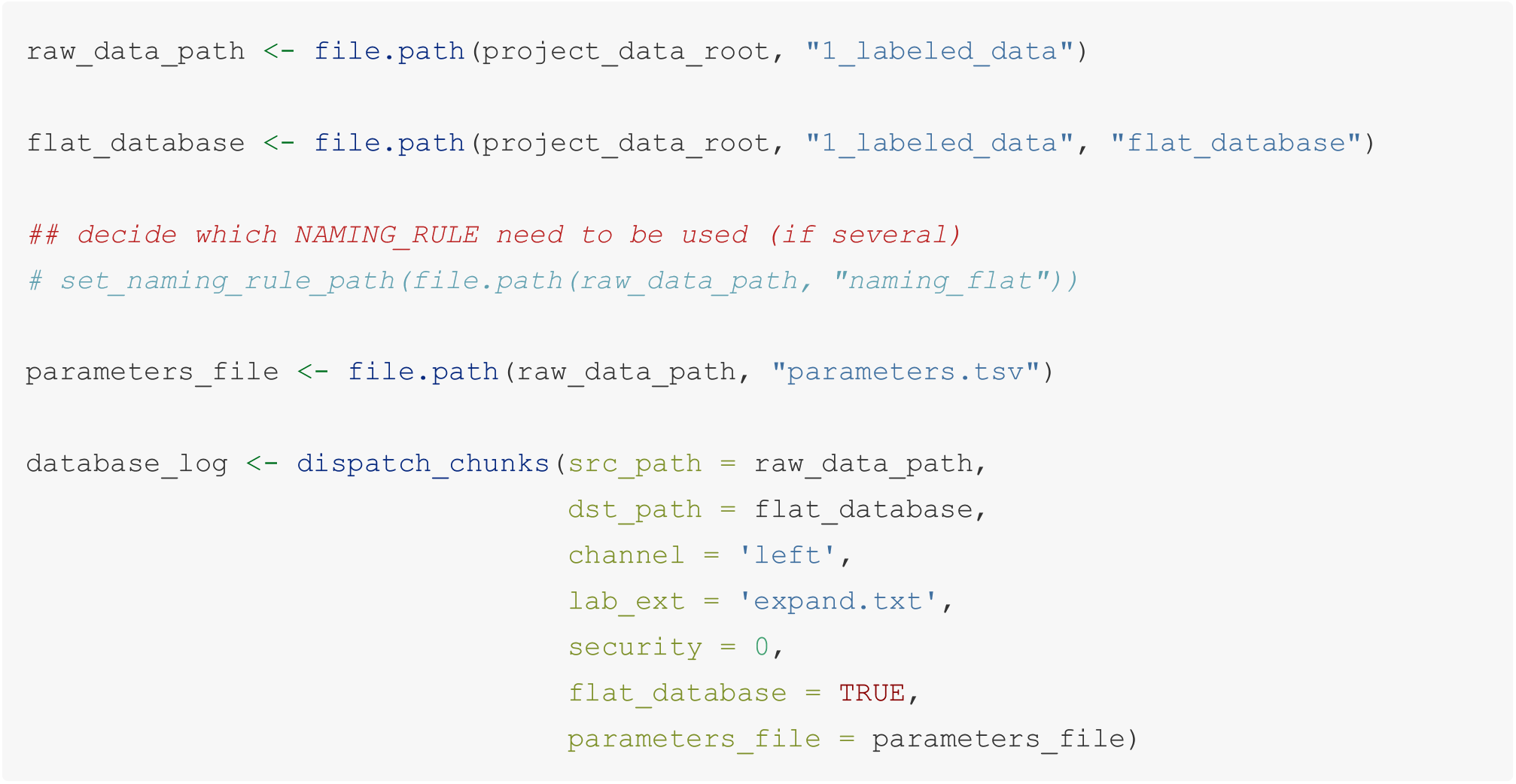

#### 1.4.2 Into a grouped database

Set up `naming_rule()` the for a grouped database.

As an example, we will dispatch the chunks per context of recording.

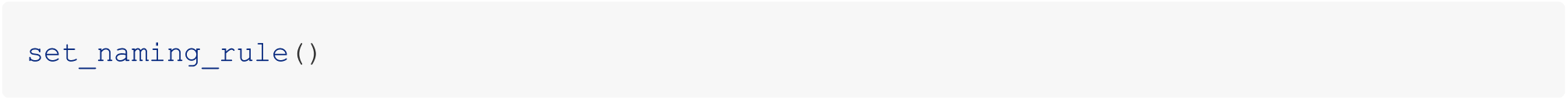

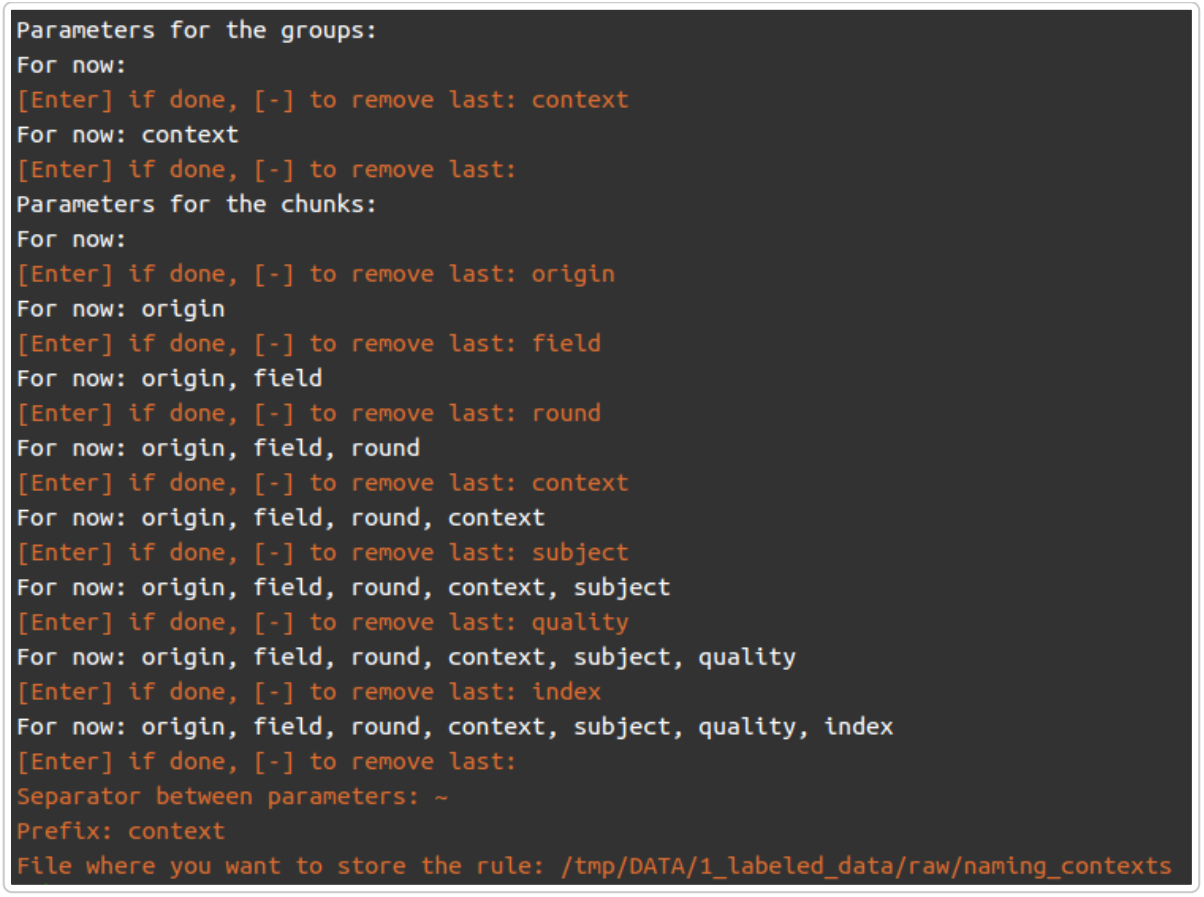

`set_naming_rule()`: user interaction for grouping into contexts

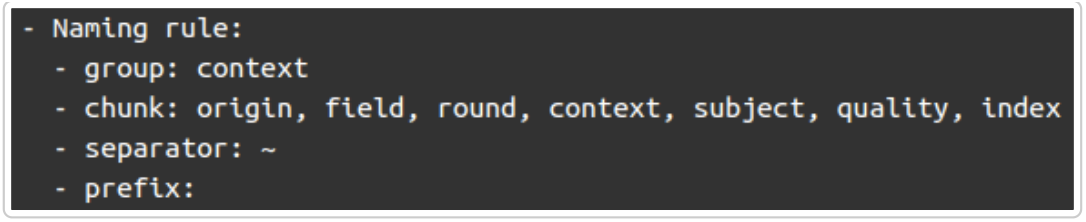

Naming rule for dispatching into `context` databases

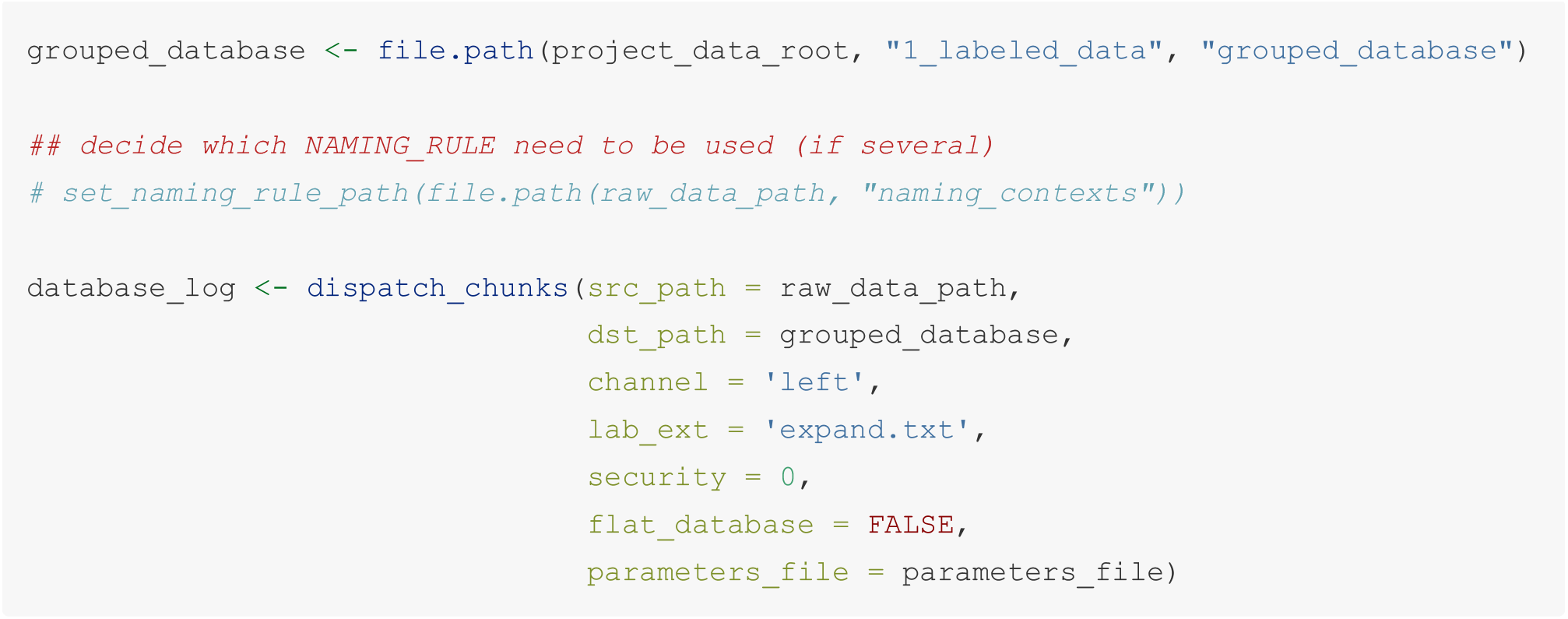

## 2 The (long) non-labeled data scenario

In this scenario, we have recordings of various durations, that most probably contain chunks, but we ignore when *a priori*.

Objective: we want to chunk these recordings in a (semi-)automated way to catch most of the chunks automatically and clean the data manually before exporting databases.

To reach this objective we need to:

- find a good set of detection settings (on slices of recordings containing some chunks)
- apply them on the entire recording to automatically chunk the recording (over all duration of recording)
- create a label track
- edit and clean up the label track if necessary
- dispatch the chunks (see first scenario)

### 2.1 Input data

A directory containing:

- a bunch of recordings
- a parameters file

What the directory looks like:

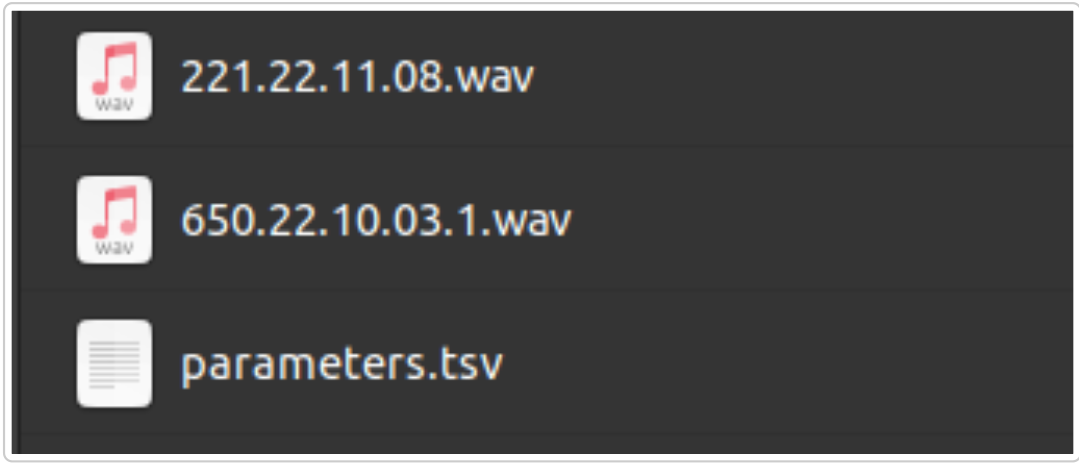

Input data directory

What the parameters file look like:

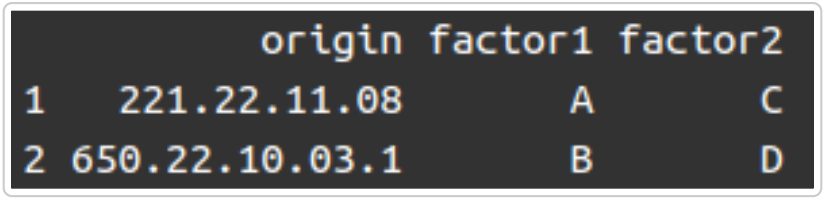

Parameters file. *Tab separated table with* .*tsv* *extension, with the column named with the* `origin` *name of the audio file, without extension*

### 2.2 Find appropriate detection settings to chunk a recording

In order to be able to chunk all recordings, we need to make sure the detection settings are relevant according to the acoustic properties of the chunks (frequency range, duration) in the recording (signal-to-noise ratio). The function `detect_chunks_interactive()` plots the spectrogram and the timer detection plot, tunes the setting and records the “best” settings in a log. However, due to its computing and memory needs, this function is better usable on short files (we advise 10 sec). Fortunately, we can generate and manage these 10 seconds slices thanks to `chop_recordings` and `dispatch`.

#### 2.2.1 Generate 10 sec slices of audio files

We define active phases of the recordings to allow the tuning of detection settings. We recommend to define these active phases such that:

- an active phase contains chunks,
- an active phase is representative of the environment of the recording (especially loudness).

We will use these active phases to create slices of recordings to find robust detection settings. What the definition of the active phases looks like in *Audacity*:

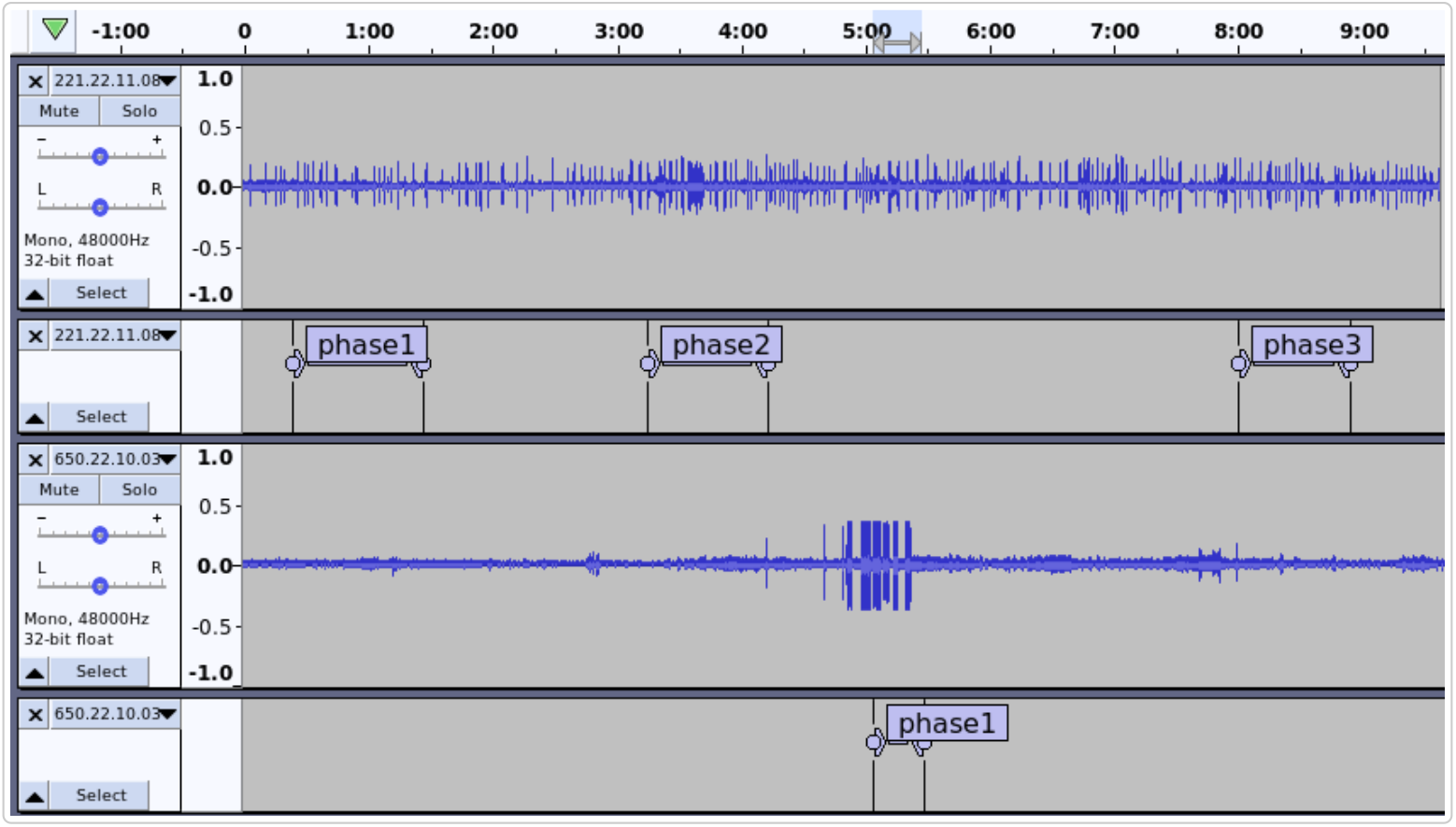

Active phases on which we will find the detection settings. Time in the recording appears in minutes.

##### a. Chop recordings into slices

We first generate the label tracks that contain the slices boundaries.

*N*.*B*. `chop_recordings()` *automatically cleans Audacity files*.

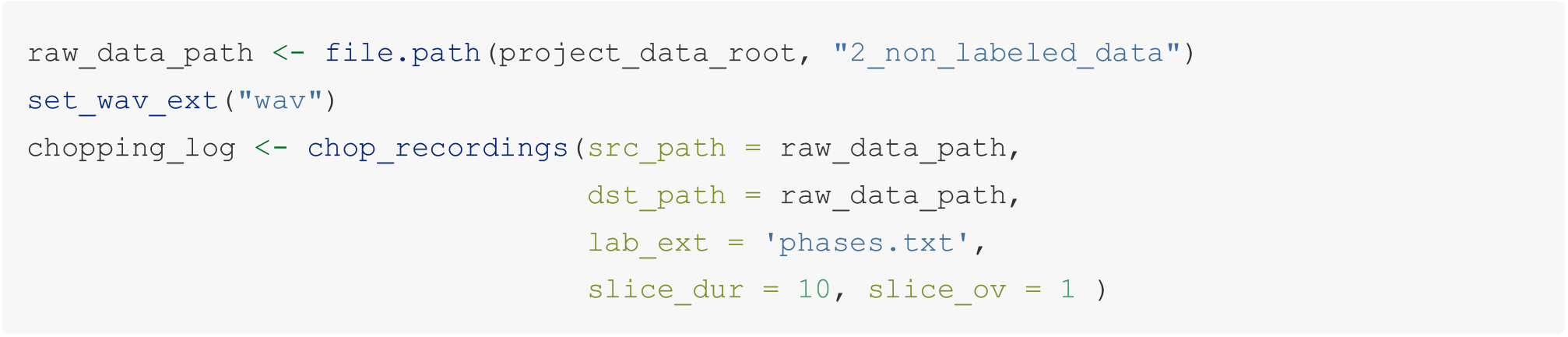

##### b. Dispatch the 10 sec slices into audio files

Slices of 10 seconds will be dispatched into a flat `tuning` database.

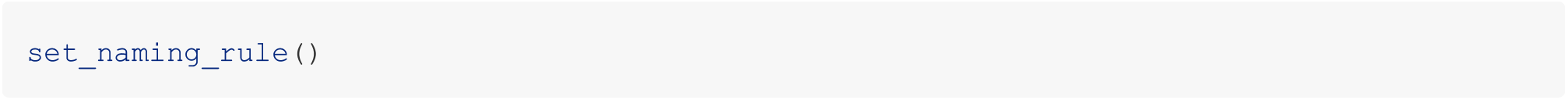

What the `naming_tuning` named `naming_rule` looks like:

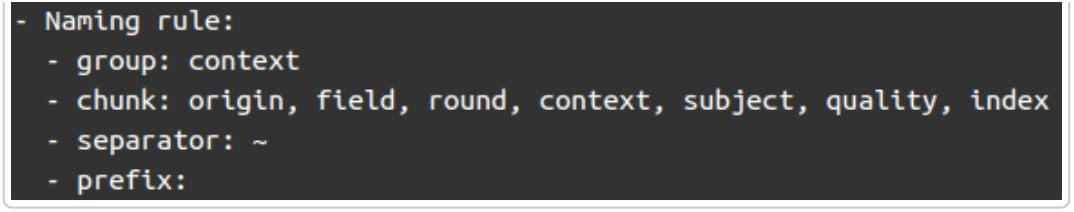

naming rule to generate tuning database

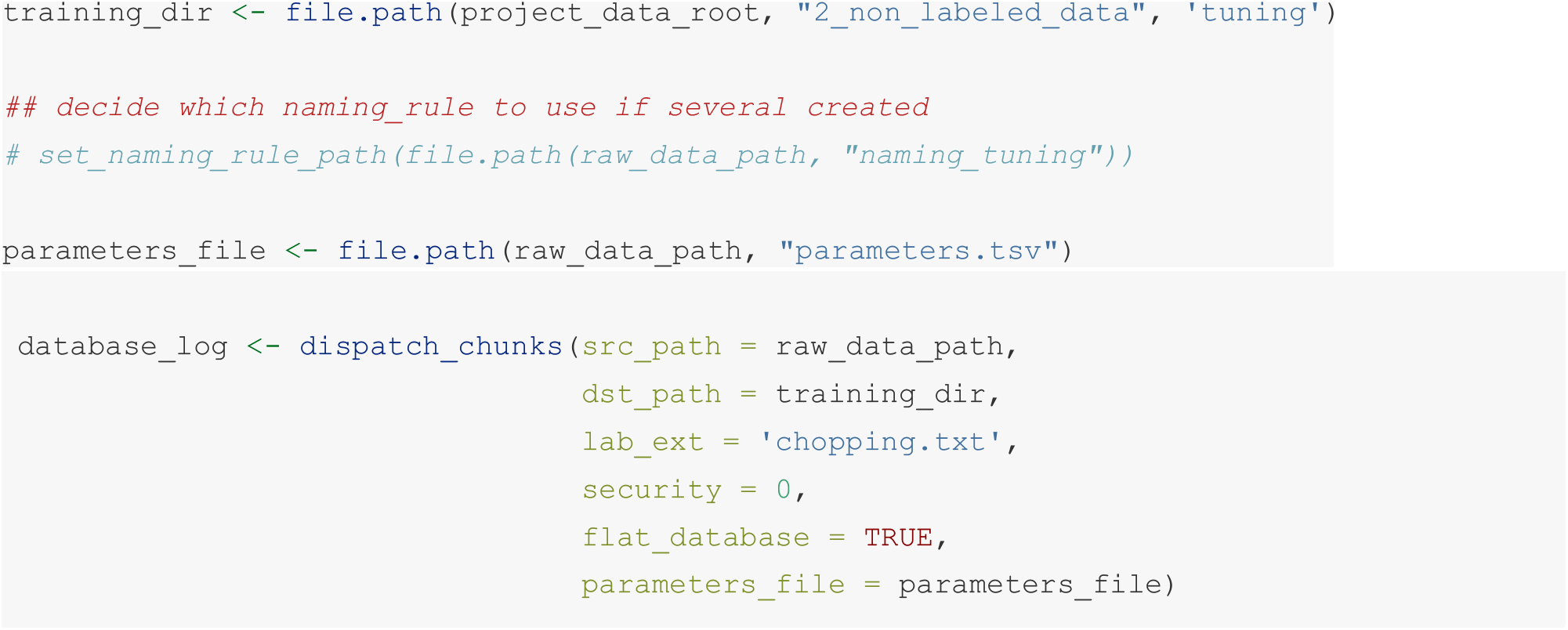

#### 2.2.2 Interactively tune detection settings

We will use this tuning database to run the `detect_chunks_interactive()` function.

The complete documentation about the function, especially for the detection settings and the forced normalisation process, is displayed through:

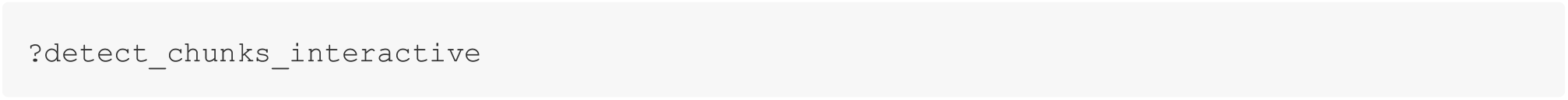

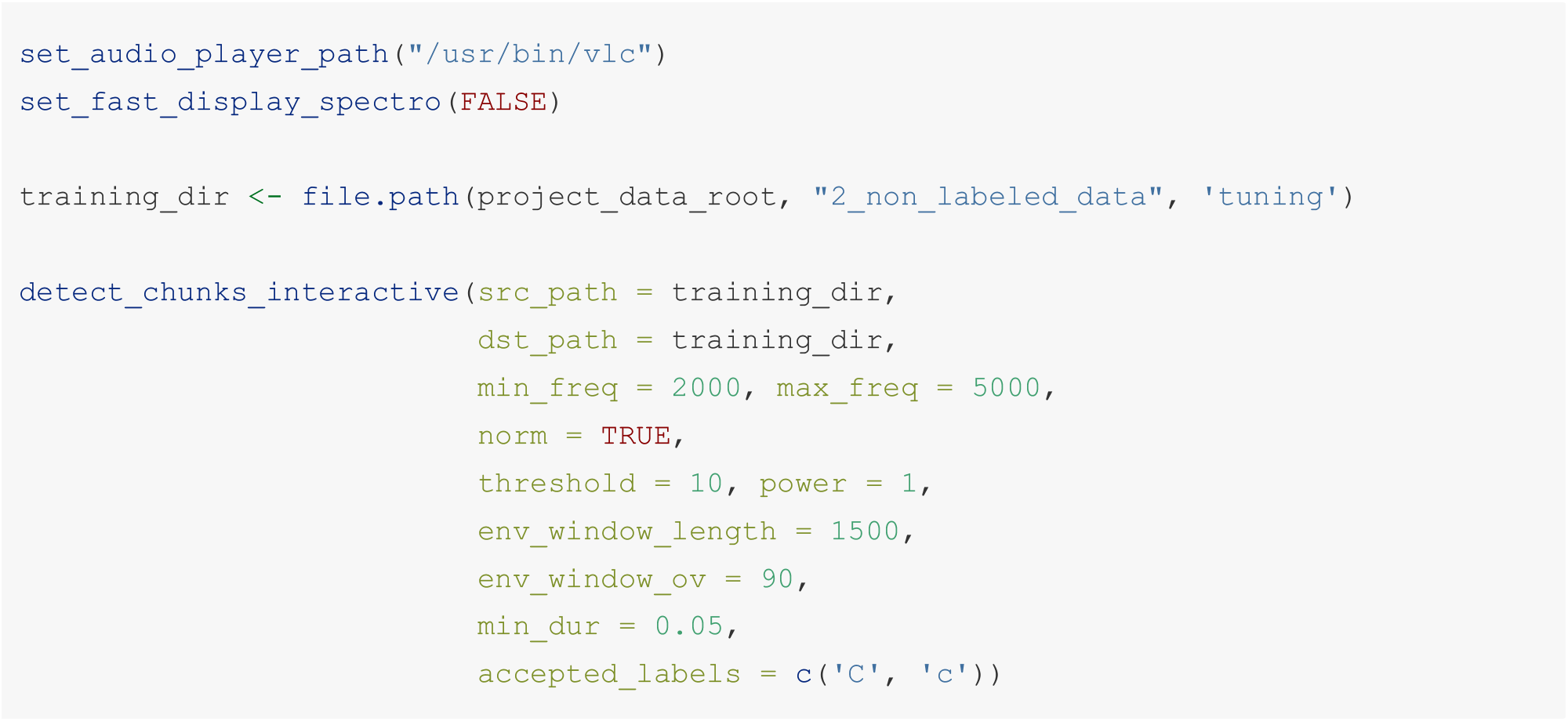

What the interface looks like in R studio:

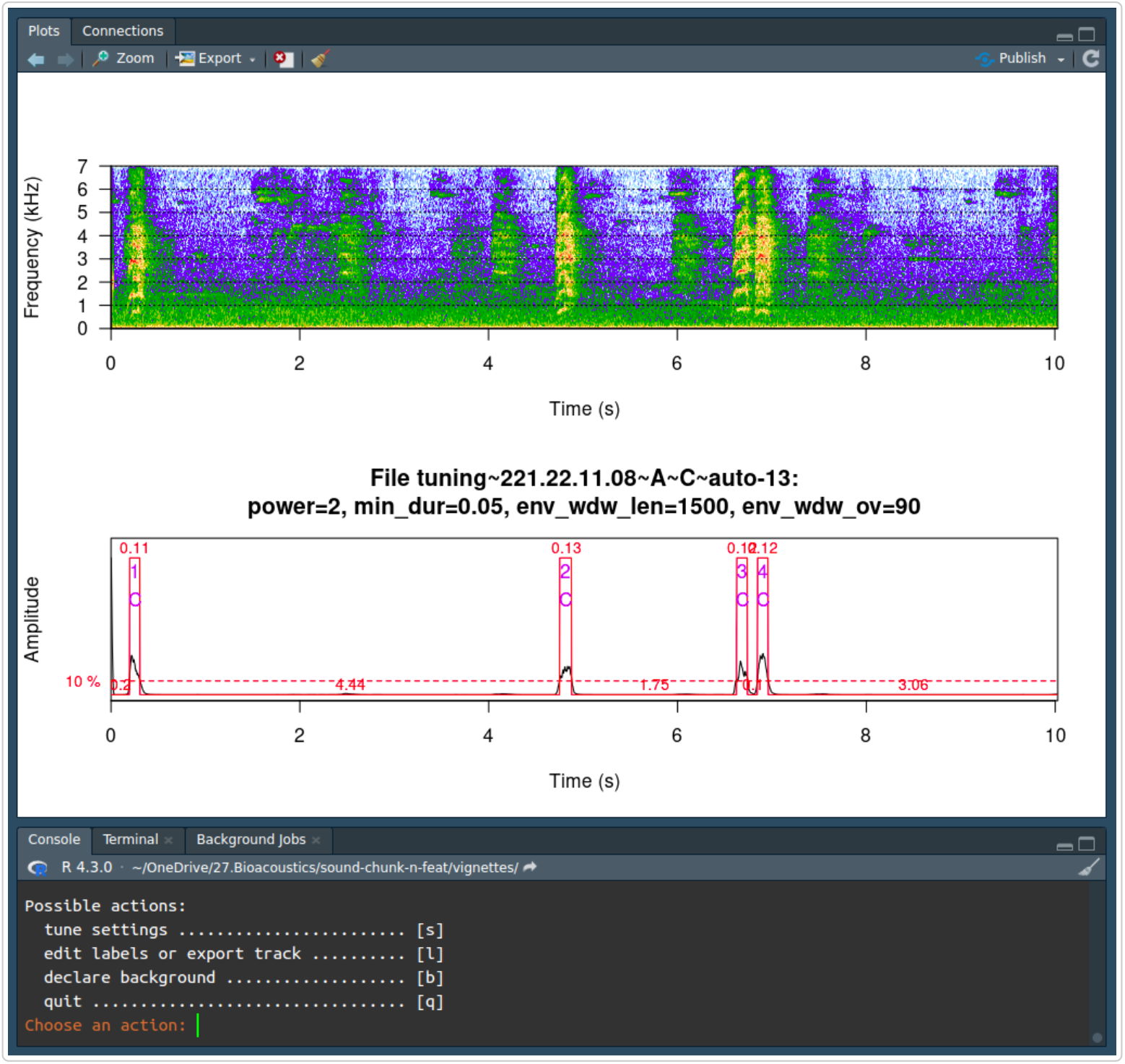

`detect_chunks_interactive()`: user interface and main menu

In the log of this function, we store the detection settings and the success rates of chunking in each 10 sec slice. This log is saved in `dst_path`. In addition, one column is used to track the normalisation value applied on all slices (if normalisation is not used, this value is NULL). This value may be used later for automated chunking.

N.B: if the `norm` argument was put to `TRUE`, the user may see an additional synthetic sound at the beginning of each slice. This is used to forced the normalisation and is discarded during the labeling process.WeWe

### 2.3 Automatically chunk recordings

The function `detect_chunks_auto` run all files of a directory and automatically chunks the recordings based on the detection settings and normalisation value.

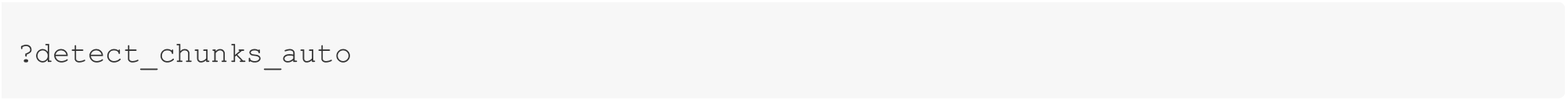

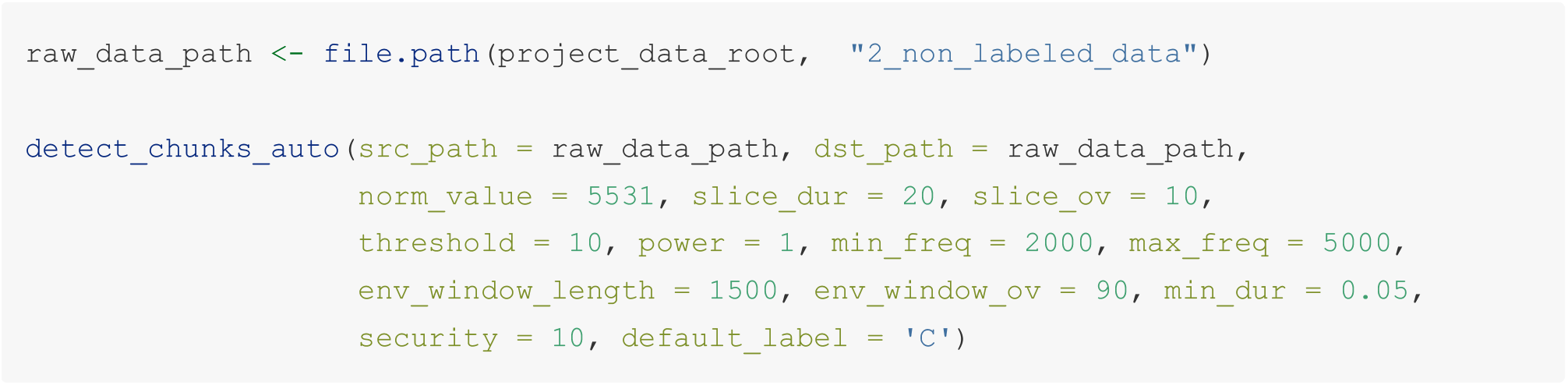

We may want to extract the calls only in specific phases of a recording, which is possible using the parameter `lab_ext`.

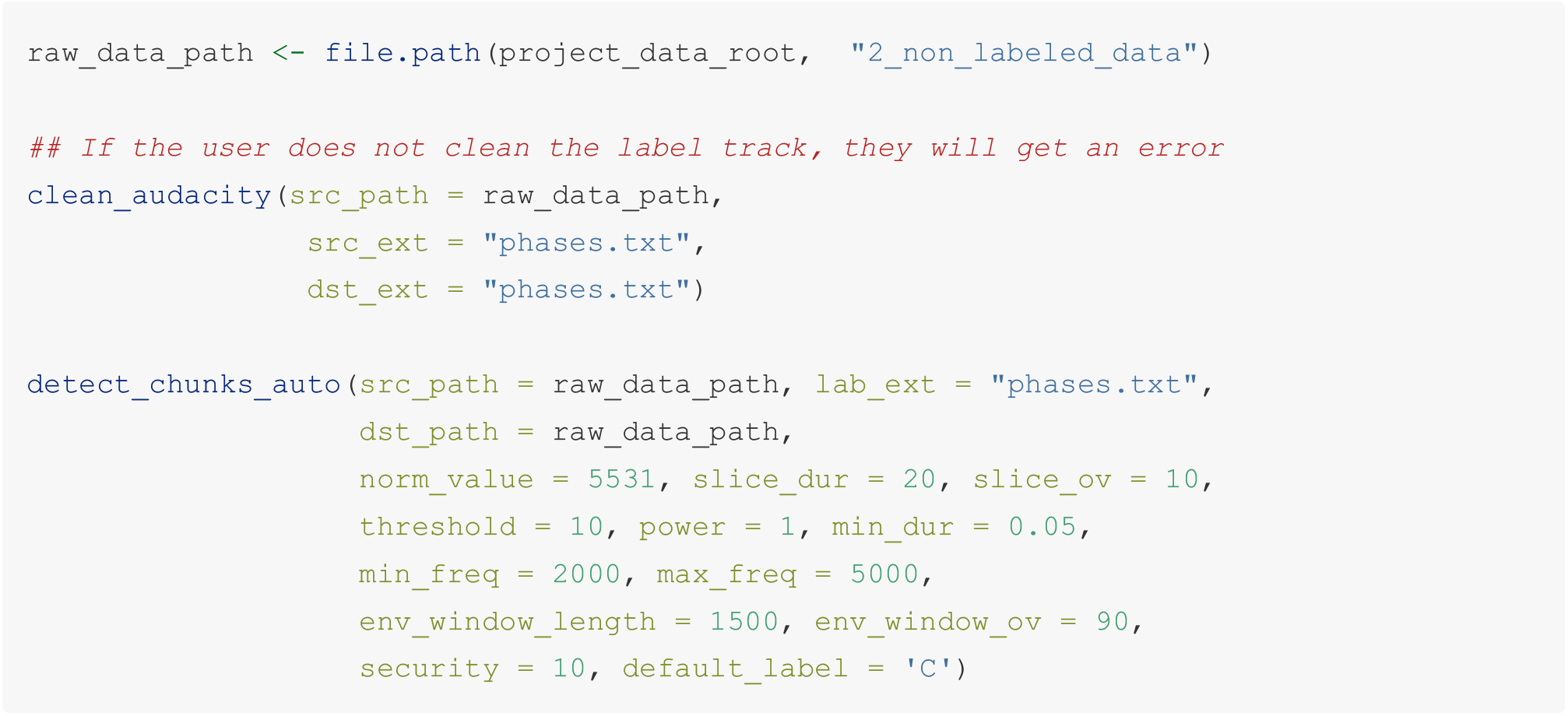

The output is one label track per recording; we may open each in *Audacity*.

What the automated label track looks like:

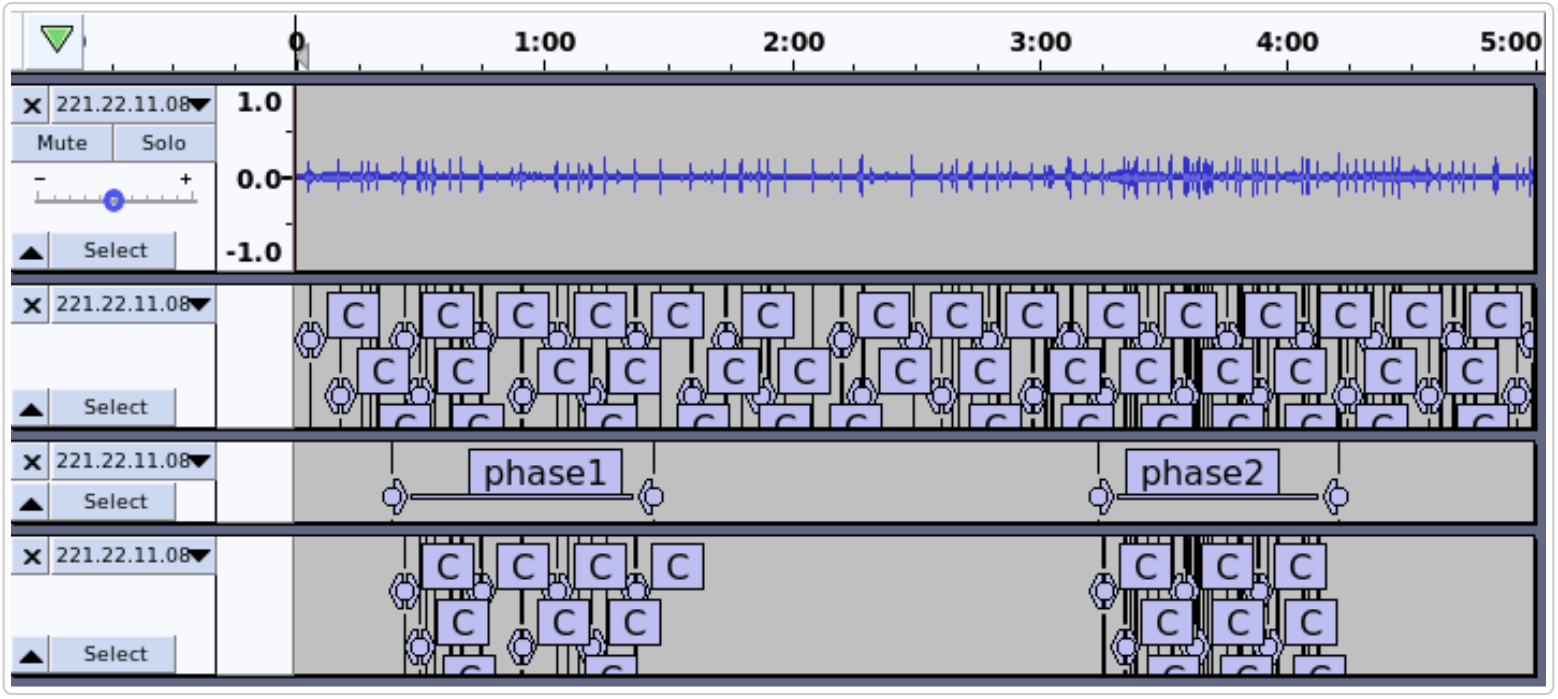

`detect_chunks_auto()`: output in Audacity, after full or partial chunking in phases.

Label tracks may be edited in *Audacity*, for example to remove potential false positive, add potential false negative, assign the quality of the chunks or add any other useful information. This label track may be re exported and used to dispatch the final chunks.

### 2.4 Dispatch the final chunks

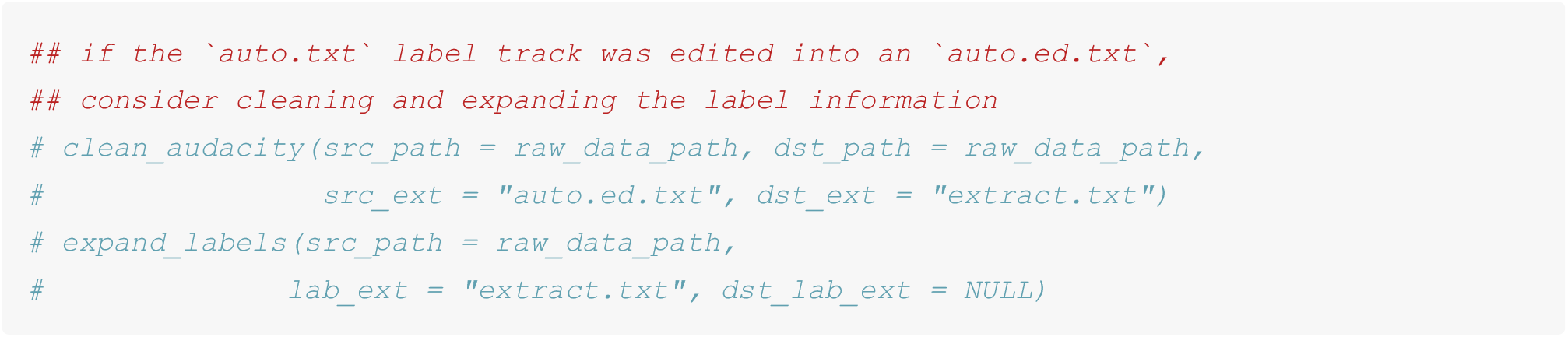

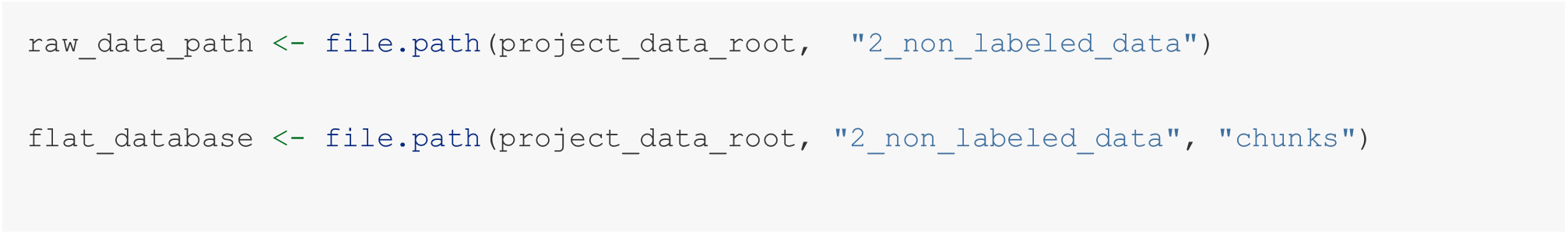

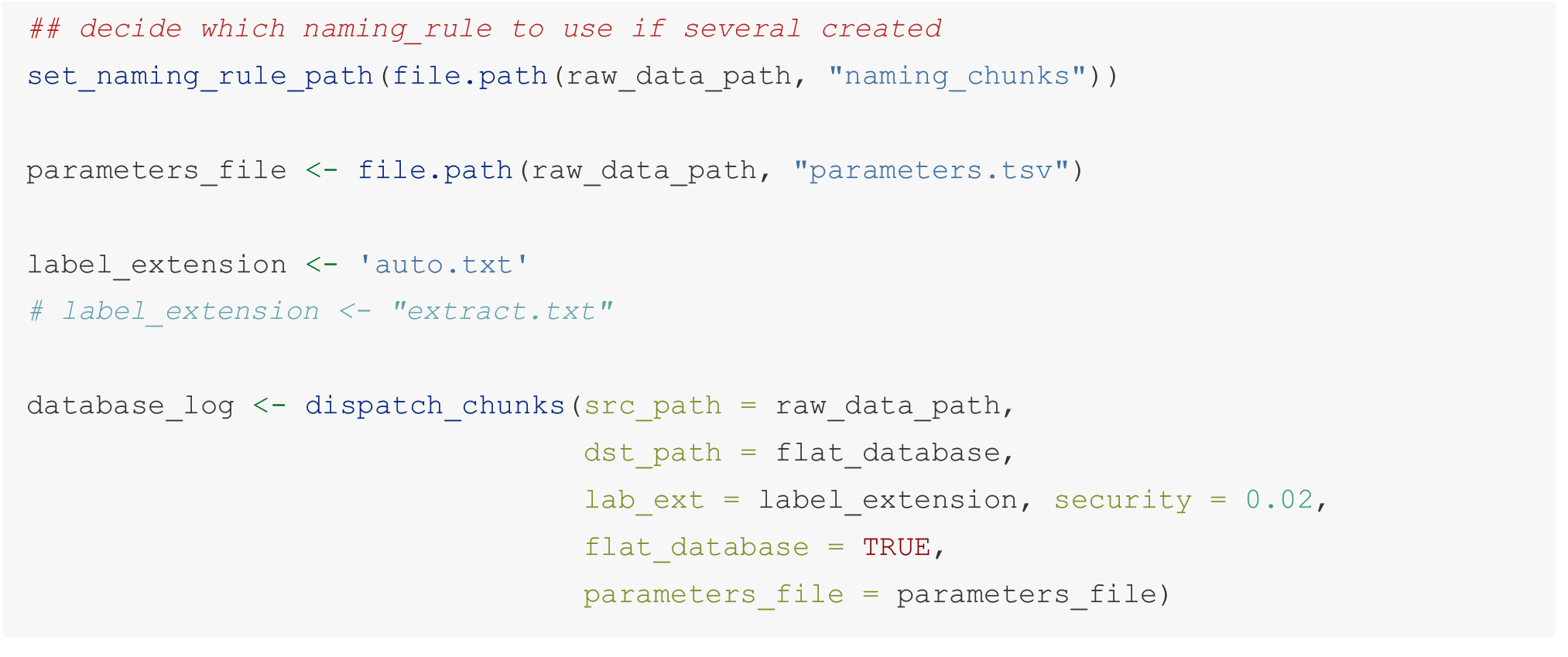

## Acknowledgements

We thank the members (and extended members) of the *Behavioural Ecology Group* who accepted to try beta versions of the functions developed in `SoundChunk`. Many thanks to Karina Stampe for sharing her parakeets recordings.

## References

“Audacity®.” 2017. https://www.audacityteam.org.

Ligges, Uwe, Sebastian Krey, Olaf Mersmann, and Sarah Schnackenberg. 2023. tuneR: Analysis of Music and Speech. https://CRAN.R-project.org/package=tuneR.

R Core Team. 2023. R: A Language and Environment for Statistical Computing. Vienna, Austria: R Foundation for Statistical Computing. https://www.R-project.org/.

Sueur, J., T. Aubin, and C. Simonis. 2008. “Seewave: A Free Modular Tool for Sound Analysis and Synthesis.” Bioacoustics 18: 213–26. https://www.tandfonline.com/doi/abs/10.1080/09524622.2008.9753600.

Villain, A. S., and P. Renaud-Goud. 2023. “Avelyne s. Villain, & Paul Renaud-Goud. (2023). SoundChunk r Package, Example Data (V1.0.0) [Data Set].” Zenodo na.: na. 10.5281/zenodo.8297684.

